# Auditory Nerve Stochasticity Impedes Auditory Category Learning: Learning: a Computational Account of the Role of Cochlear Nucleus and Inferior Colliculus in Stabilising Auditory Nerve Firing

**DOI:** 10.1101/059428

**Authors:** Irina Higgins, Simon Stringer, Jan Schnupp

**Affiliations:** Department of Experimental Psychology, University of Oxford, Oxford, England; Department of Physiology, Anatomy and Genetics (DPAG), University of Oxford, Oxford, England

## Abstract

It is well known that auditory nerve (AN) fibers overcome bandwidth limitations through the “volley principle”, a form of multiplexing. What is less well known is that the volley principle introduces a degree of unpredictability into AN neural firing patterns which makes even simple stimulus categorization tasks difficult. We use a physiologically grounded, unsupervised spiking neural network model of the auditory brain with STDP learning to demonstrate that plastic auditory cortex is unable to learn even simple auditory object categories when exposed to the raw AN firing input without subcortical preprocessing. We then demonstrate the importance of non-plastic subcortical preprocessing within the cochlear nucleus (CN) and the inferior colliculus (IC) for stabilising and denoising AN responses. Such preprocessing enables the plastic auditory cortex to learn efficient robust representations of the auditory object categories. The biological realism of our model makes it suitable for generating neurophysiologically testable hypotheses.

## Introduction

The hierarchy of the auditory brain is complex, with numerous interconnected subcortical and cortical areas. While a wealth of neural response data has been collected from the auditory brain [1–3], the role of the computations performed within these areas and the mechanism by which the sensory features of auditory objects are transformed into higher-order representations of object category identities are yet unknown [4]. How does the auditory brain learn robust auditory categories, such as phoneme identities, despite the large acoustical variability exhibited by the raw auditory waves representing the different auditory object exemplars belonging to a single category? How does it cope once this variability is further amplified by the spike time stochasticity inherent to the auditory nerve (AN) when the sounds are encoded into neuronal discharge patterns within the inner ear?

One of the well accepted theories explaining the information encoding operation of the AN is the so called “volley principle” [5]. It states that groups of AN fibers with a similar frequency preference tend to phase-lock to different randomly selected peaks of a simple sinusoidal sound wave when the frequency of the sinusoid is higher than the maximal frequency of firing of the AN cells. This allows the AN to overcome its bandwidth limitations and represent high frequencies of sound through the combined frequency of firing within groups of AN cells. It has not been considered before, however, that the information encoding benefits of the volley principle may come at a cost. Here we suggest that this cost is the addition of the so called “spatial jitter” to the AN firing.

It is useful to think of the variability in AN discharge patterns as a combination of “temporal” and “spatial jitter”. Temporal jitter arises when the AN fiber propensity to phase lock to temporal features of the stimulus is degraded to a greater or lesser extent by poisson-like noise in the nerve fibers and refractoriness [6]. “Spatial jitter” refers to the fact that neighbouring AN fibers have almost identical tuning properties so that an action potential that might be expected at a particular fiber at a particular time may be observed in one of the neighbouring fibers [5]. In this paper we argue that space and time jitter obscure the similarities between the AN spike rasters in response to different presentations of auditory stimuli belonging to the same class, thus impeding auditory object category learning.

The reason why we believe that excessive amount of jitter in the AN can impair auditory object category learning in the auditory cortex is the following. Previous simulation work has demonstrated that one way category learning can arise in competitive feedforward neural architectures characteristic of the cortex is through the “continuous transformation” (CT) learning mechanism [7,8]. CT learning is a biologically plausible mechanism based on Hebbian learning, which operates on the assumption that highly similar, overlapping input patterns are more likely to be different exemplars of the same stimulus class. CT learning then binds these similar input patterns together onto the same subset of higher stage neurons, which, thereby, learn to be selective and informative about their learnt preferred stimulus class. The CT learning principle is a biologically plausible mechanism for learning object transformation orbits as described by [9]. CT learning breaks when the similarity between the nearest neighbour exemplars within a stimulus class become approximately equal to the similarity between the nearest neighbour exemplars in different stimulus classes. A more detailed description of CT learning is provided in the Materials and Methods section.

In this paper we argue that the additional spike time variability introduced in the AN input representations of the different exemplars belonging to a single auditory object class break CT learning. We show this by training a simple biologically realistic feedforward spiking neural network model of the auditory cortex with spike timing dependent plasticity (STDP) learning [10] to perform simple categorisation of two synthesised vowel classes using raw AN firing as input (AN-A1 model shown in Fig. 1B). We show that such a model is unable to solve this easy categorisation task because the reproducibility of AN firing patterns for similar stimuli necessary for CT learning to operate is disrupted by the multiplexing effects of the volley principle in the AN.

**Fig 1.**
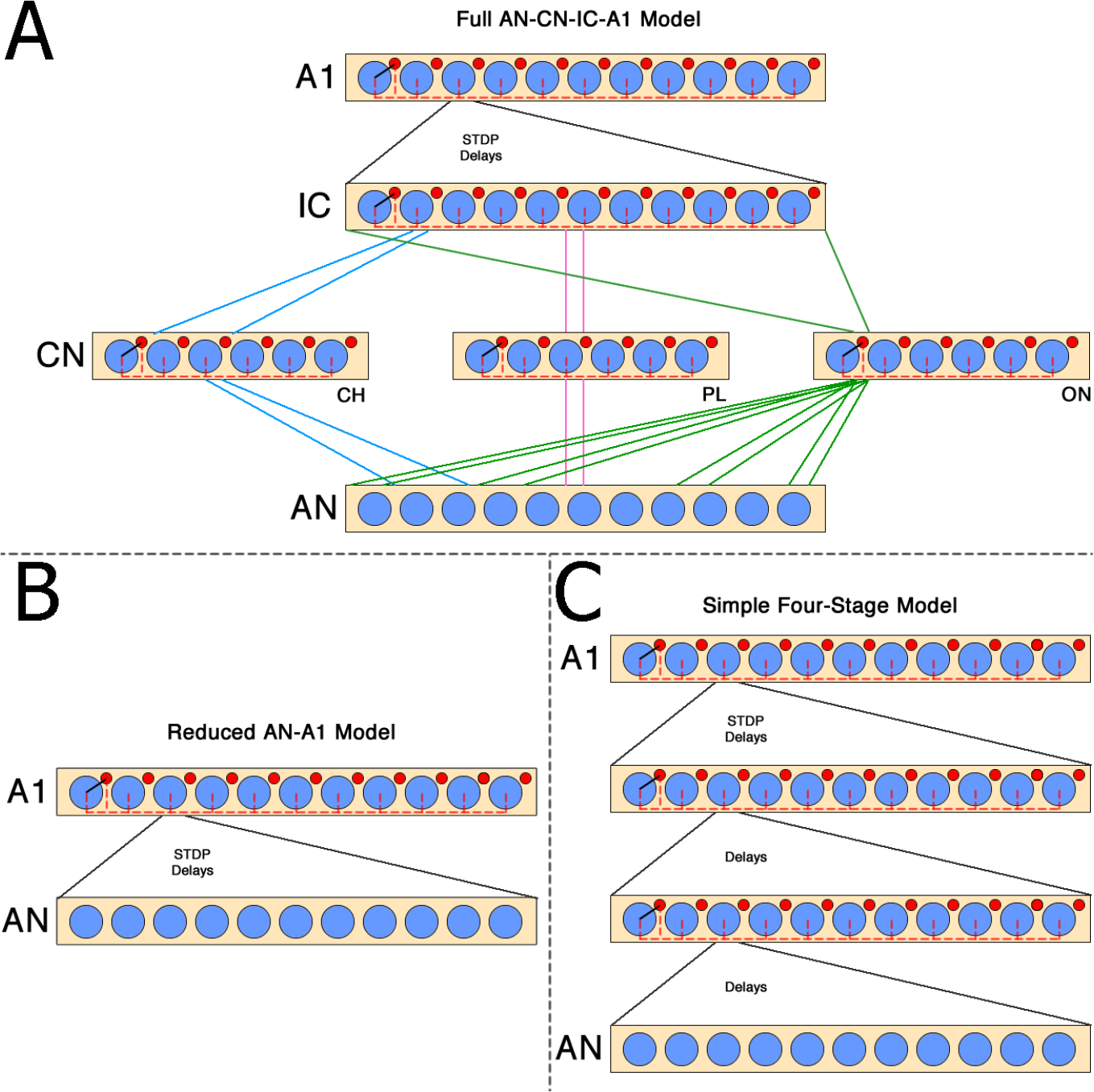
Schematic representation of the full AN-CN-IC-A1 (**A**), the reduced AN-A1 (**B**) and the simple four-stage (**C**) models of the auditory brain. Blue circles represent excitatory (E) and red circles represent inhibitory (I) neurons. The connectivity within each stage of the models is demonstrated using one excitatory cell as an example: **E→I** connection is shown in black, **I→E** connections are shown in red. feedforward connections between the last two stages of each model are modifiable through STDP learning. AN-auditory nerve; CN-cochlear nucleus with three subpopulations of cells: chopper (CH), primary-like (PL) and onset (ON), each exhibiting different response patterns by virtue of their distinct connectivity; IC-inferior colliculus; A1-primary auditory cortex.

This observation has led us to believe that an extra preprocessing stage was necessary between the AN and the plastic A1 in order to reduce the jitter (noise) found in the temporal and spatial distribution of AN spikes in response to the different exemplars of the same auditory stimulus class. This reduction in jitter would be necessary to enable the plastic auditory cortex to learn representations of auditory categories through CT learning. We hypothesised that this preprocessing could happen in the intermediate subcortical stages of processing in the auditory brain, such as CN and IC, whereby the essential contribution of the precise microarchitecture and connectivity of the CN and IC would be to help de-jitter and stabilise the AN firing patterns, thereby enabling the plastic cortical area A1 to develop informative representations of vowel categories through CT learning.

We tested our hypothesis by comparing the performance of a biologically realistic four stage hierarchical feedforward spiking neural network model of the auditory brain incorporating both subcortical (AN, CN, IC) and cortical (A1) stages (full AN-CN-IC-A1 model shown in Fig. 1A) to the performance of two models that either omitted areas CN and IC (reduced AN-A1 model shown in Fig. 1B), or had the same number of processing stages as the full AN-CN-IC-A1 model, but lacked the precise CN and IC microarchitecture and connectivity (simple four-stage model shown in Fig. 1C). Our simulations demonstrated that both the reduced AN-A1 and simple four-stage models significantly underperformed the full AN-CN-IC-A1 model on the two vowel classification task.

The contributions of this work are two-fold: 1) it shows how simple, local synaptic learning rules can support unsupervised auditory category learning if, and only if, stochastic noise introduced when sounds are encoded at the auditory nerve is dealt with by auditory brainstem processes; 2) it provides a quantitative theoretical framework which explains the diverse physiological response properties of identified cell classes in the ventral cochlear nucleus and generates neurophysiologically testable hypotheses for the essential role of the non-plastic CN and IC as the AN jitter removal stages of the auditory brain.

## Results

### Quantifying Spike Jitter in the Auditory Nerve

In this paper we postulate that the reproducibility of AN firing patterns for similar stimuli necessary for CT learning to operate is disrupted by the multiplexing effects of the volley principle in the AN. This is supported by a quantitative analysis of the similarity/dissimilarity between AN firing patterns in response to vowel stimuli belonging either to the same or different synthesised vowel classes (see Materials and Methods for calculation details). Indeed, it was found that the AN spike rasters for repeat presentations of the same exemplar of a vowel or of different exemplars of the same vowel category were as dissimilar to each other as the AN responses to the vowels from different vowel categories (see the AN scores in Tbl. 1).

**Table 1.** Similarity measure scores between the AN and IC spike rasters in response to: (i) different presentations of the same exemplar of a stimulus (Same Exemplar Index), (ii) different exemplars of the same stimulus class (Different Exemplars Index), and different stimulus classes (Different Categories Index). Scores vary between 0 and 1, with higher scores indicating higher levels of similarity and consequently low levels of jitter.

|  | /i:/ |  | /a/ |  | /i:/ and /a/ |  |
| --- | --- | --- | --- | --- | --- | --- |
|  | AN | IC | AN | IC | AN | IC |
| Same Exemplar Index | 0.45 | 0.9 | 0.57 | 1 | - | - |
| Different Exemplars Index | 0.52 | 0.91 | 0.63 | 1 | - | - |
| Different Categories Index | - | - | - | - | 0.42 | 0.67 |

### Reduced AN-A1 Auditory Brain Model

We begin by presenting simulation results from the reduced AN-A1 spiking neural network model of the auditory brain shown in Figure 1B, in which the intermediate CN and IC stages were omitted (see Materials and Methods for model architecture details). The input stage of the AN-A1 model is a highly biologically realistic AN model by [11], and the output stage is a loose and simplified approximation of the A1 in the real brain.

We tested the ability of the AN-A1 model to learn robust representations of auditory categories using a controlled yet challenging task, whereby twelve different exemplars of each of two classes of vowels, /i:/ and /a/, were synthesised and presented to the network (Fig. 2) (see Materials and Methods). The biologically plausible unsupervised CT learning mechanism implemented through STDP within the AN→A1 afferent connections was expected to enable the model to learn the two vowel categories (see Materials and Methods for an overview of CT learning). In particular, we investigated whether localist representations of auditory categories emerged, whereby individual neurons would learn to respond selectively to all exemplars of just one preferred stimulus class [12].

**Fig 2.**
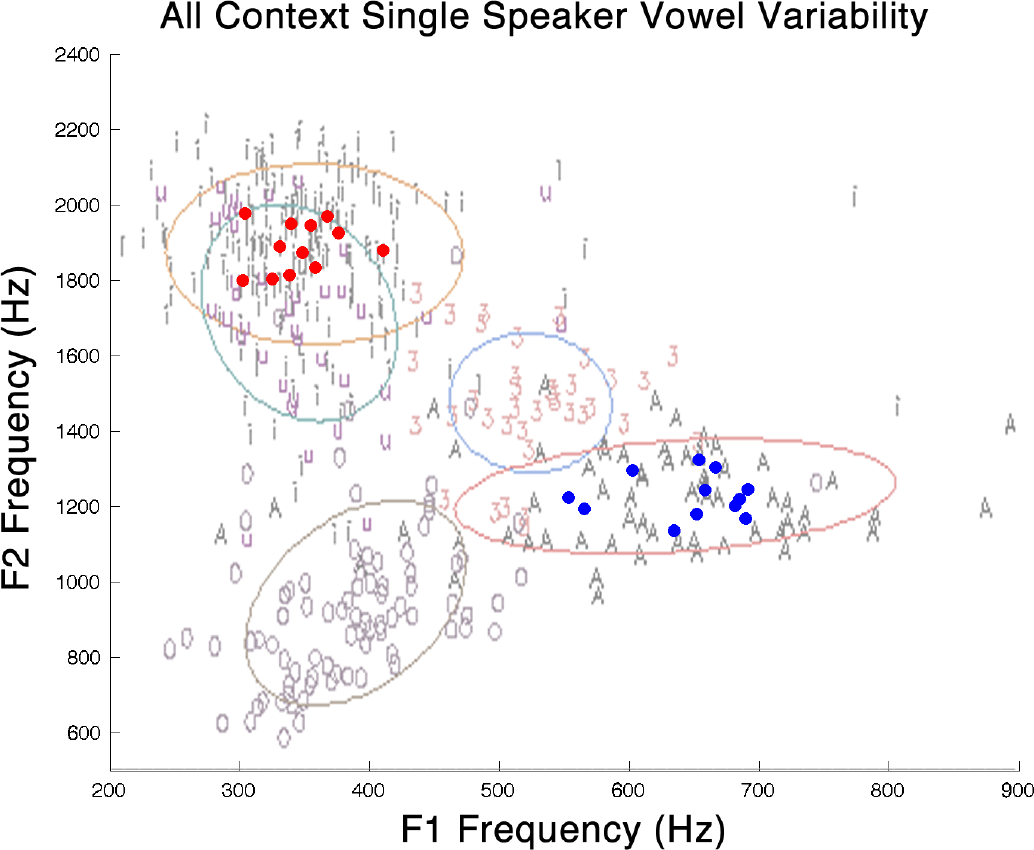
Schematic representation of twelve transforms of two synthesised vowels (/a/-blue, /i:/-red) projected onto the two-dimensional plane defined by the first two formants of the vowels. Each transform was generated by randomly sampling three formant frequencies from a uniform 200 Hz distribution centered around the respective average values reported by [13] for male speakers. It can be seen that the generated vowel transforms are in line with the vowel distribution clouds produced from natural speech of a single speaker [14]. All transforms were checked by human subjects to ensure that they were recognisable as either an /a/ or an /i:/. The ellipses approximate the 70% within-speaker variability boundary for a particular phoneme class.

The ability of the AN-A1 model to learn robust vowel categories depends on how it is parameterized. A hyper-parameter search using a grid heuristic was, therefore, conducted. Mutual information between the stimuli and the responses of singles cells within the output A1 stage of the model was used to evaluate the performance of the AN-A1 model on the vowel categorization task (see Materials and Methods). It was assumed that the performance of the network changed gradually and continuously as a function of its hyper-parameters, since learning in the real brain has to be robust to mild variations in biological parameters. It was, therefore, expected that the best model performance found through the grid parameter search would be a good approximation of the true maximal model performance. The detailed description of the parameter search can be found in Supplemental Materials. The following parameters were found to result in the best AN-A1 model performance: LTP constant (*α_p_*) = 0.05; LTD constant (*α_d_*) = −0.02; STDP time constants (*τ_p_/τ_d_*) = 15/25 ms; initialisation magnitude of AN→ A1 connections 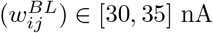; level of inhibition in the A1 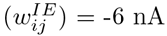.

The performance of the best AN-A1 model found through the parameter search is shown in Fig. 3 (solid dark blue line). The average information about the vowel class identity among the top ten most informative A1 cells was 0.21 bits and the maximum A1 information was 0.57 bits out of the theoretical maximum of 1 bit. This is not enough to achieve good vowel recognition performance, even though a certain amount of useful learning did occur in the reduced AN-A1 model as evidenced by more A1 information after training than before training, and more information in the A1 compared to the AN input (Fig. 3, dotted dark blue and solid red lines respectively).

**Fig 3.**
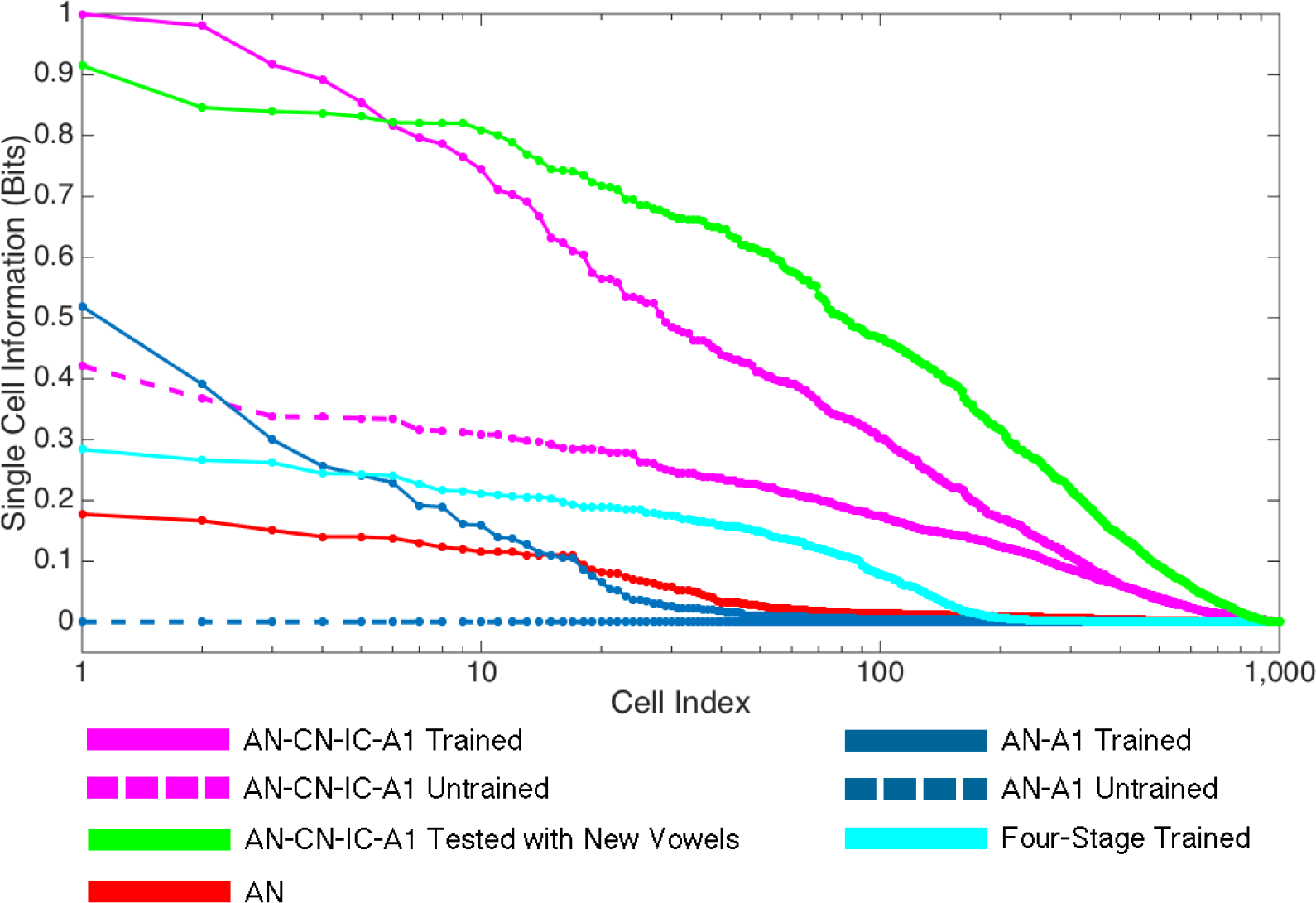
Single cell information carried by cells in a specified model neural area during the vowel classification task. The cells are ordered along the abscissa by their informativeness. Maximum theoretical entropy for the task is 1 bit. It can be seen that the output A1 neurons of the full AN-CN-IC-A1 spiking neural network model of the auditory brain after training carry more information about the two vowel classes than the input auditory nerve (AN) fibers, or the A1 cells of the reduced AN-A1 model, simple four-stage model, or any of the models before training.

### Removing Auditory Nerve Jitter

The reduced AN-A1 model was unable to learn the identities of the two vowel classes through unsupervised CT learning implemented through STDP within the plastic AN→A1 connections. Successful CT learning relies on the discovery of correlations, or “overlap”, in the neural representations of stimuli that belong to the same “object” or stimulus class. We attribute the failure of the A1 neurons in the reduced model to discover stimulus classes to the fact that the highly biologically realistic AN input to the model contains large amounts of physiological noise or “space” and “time” jitter in the spike times, which obscure the similarities between the AN spike rasters in response to different stimuli belonging to the same vowel class. Since such similarities are necessary for CT learning to operate, the output A1 stage of the reduced AN-A1 model was unable to learn robust representations of the two vowel classes directly from the AN input.

Reducing time and space jitter in AN response spike rasters should aid unsupervised learning in the auditory brain, and it can be achieved through the following mechanisms: 1) information from a number of AN fibers with similar characteristic frequencies (CFs) is integrated in order to remove space jitter; and 2) AN spike trains for different cells are synchronised, whereby spikes are re-aligned to occur at set points in time rather than anywhere in continuous time, thus removing time jitter.

We consider space and time jitter removal to be one of the key roles of the subcortical areas CN and IC, whereby jitter reduction is initiated in the CN and completed within the IC, as convergent inputs from different subpopulations of the CN are integrated in such a way which facilitates effective stimulus classification by CT-like learning mechanisms in subsequent stages, such as A1. We envisage the following processes: 1) chopper (CH) cells within the CN remove space jitter; 2) onset (ON) cells within the CN remove time jitter; 3) the IC produces spike rasters with reduced jitter in both space and time by combining the afferent activity from the cochlear nucleus CH and ON cells.

#### Space Jitter Removal

CH neurons in the CN are suitable for the space jitter removal task due to their afferent connectivity patterns from the AN. Each CH cell receives a small number of afferent connections from AN neurons with similar CFs [15]. The incoming signals are integrated to produce regular spike trains.

In the full AN-CN-IC-A1 model shown in Fig. 1A, a CH subpopulation was simulated by adding 1000 Class 1 neurons by [16] with Gaussian topological connectivity from the AN, whereby each CH cell received afferents from a tonotopic region of the AN. A hyper-parameter search was conducted to maximise the space jitter removal ability of CH neurons (see Supplemental Materials), and the following parameter values were found to be optimal: Gaussian variance of the AN→CH afferent connectivity (*σ*) = 26 cells; magnitude of the AN→CH afferent connections 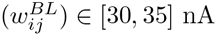; within-CH inhibition 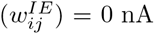. The discharge properties of the optimised CH cells corresponded closely to those reported experimentally for biological CH neurons (Fig. 4, right column).

**Fig 4.**
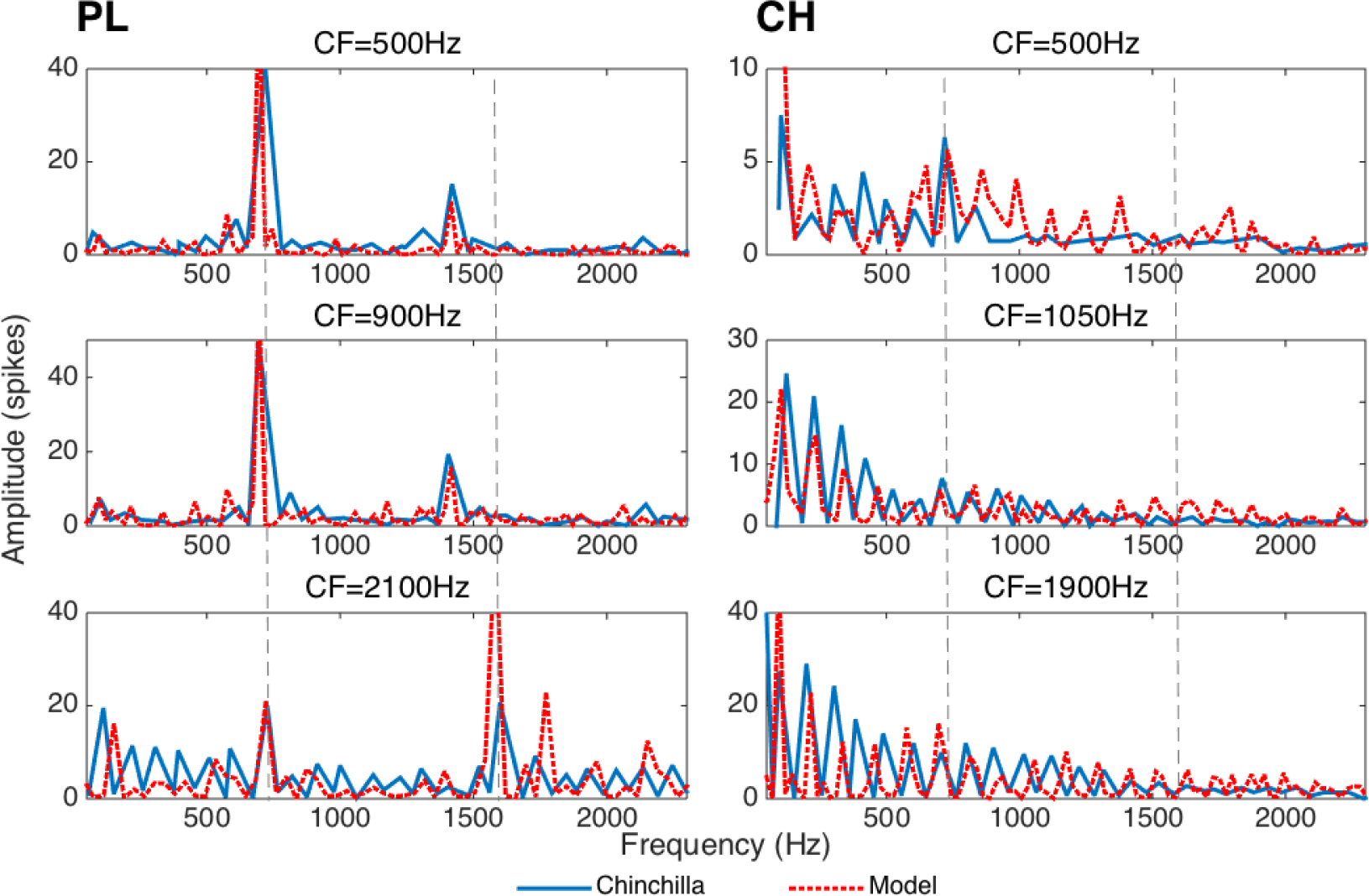
Spectra (computed as Fast Fourier Transforms of period histograms) of primary-like (PL) (left column) and chopper (CH) (right column) cochlear nucleus neuron responses to a synthetic vowel /a/ generated using the Klatt synthesiser [17]. The ordinate represents the level of phase-locking to the stimulus at frequencies shown along the abscissa. Dotted lines show the positions of the vowel formant frequencies *F*_1_ and *F*_2_. Data from chinchilla CN fibers reproduced from [2] is shown in solid blue. Data collected from the corresponding model CN fibers is shown in dashed red. Similarity between the real and model fibers’ response properties suggests that the model’s performance is comparable to the neurophysiological data.

#### Time Jitter Removal

Time jitter removal is thought to be facilitated by ON neurons in the CN. ON cells are relatively rare, constituting approximately 10% of the ventral CN [18]. They have been estimated to each receive connections from up to 65 AN fibers across a wide stretch of the cochlea, which results in broadly frequency tuned response properties [18]. These cells are characterised by fast membrane time constants, which makes them very leaky, with high spike thresholds. Consequently, ON cells require strong synchronisation from many AN fibers with a wide range of CFs in order to produce a discharge [19]. The cross frequency coincidence detection inherent to the ON cells makes them able to phase-lock to the fundamental frequency (*F*_0_) of vowels, as supported by neurophysiological evidence [20].

We propose that the interplay between the converging ON and CH cell inputs to the IC can reduce jitter in the neural representations of vocalisation sounds. Since ON cells synchronise to the stimulus *F*_0_, they can introduce regularly spaced afferent input to the IC. Such subthreshold afferent input would prime the postsynaptic IC cells to discharge at times corresponding to the cycles of stimulus *F*_0_. If IC cells also receive input from CH cells, then ON afferents will help synchronise CH inputs within the IC by increasing the likelihood of the IC cells firing at the beginning of each *F*_0_ cycle. This is similar to the encoding hypothesis described in [21].

In the full AN-CN-IC-A1 model, a population of ON cells was simulated using 100 Class 1 neurons by [16] sparsely connected to the AN. A hyper-parameter search was conducted to maximise the ability of ON neurons to synchronise to the *F*_0_ of the stimuli (see Supplemental Materials), and the following parameter values were found to be optimal: AN→ON afferent connection weight magnitudes 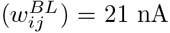; sparseness of AN-ON connectivity = 0.46 (54% of all possible AN-ON connections are non-zero); within-ON inhibition magnitude 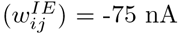.

### Full AN-CN-IC-A1 Auditory Brain Model

The full AN-CN-IC-A1 model of the auditory brain was constructed as shown in Fig. 1A to test whether the addition of the subcortical stages corresponding to the CN and IC would remove space and time jitter contained within the input AN firing rasters as described above, and thus enable the output plastic cortical stage A1 to learn invariant representations of the two vowel categories, /i:/ and /a/ (see Materials and Methods for details of the model architecture). Similarly to the reduced AN-A1 model, the output stage of the full AN-CN-IC-A1 model is a loose and simplified approximation of the A1 in the real brain.

In the brain sub-populations of the CN do not necessarily synapse on the IC directly. Instead, they pass through a number of nuclei within the superior olivary complex (SOC). The nature of processing done within the SOC in terms of auditory object recognition (rather than sound localisation), however, is unclear. The information from the different CN sub-populations does converge in the IC eventually, and for the purposes of the current argument we model this convergence as direct. The same simplified connectivity pattern (direct CN-IC projections) was implemented by [22] for their model of the subcortical auditory brain.

Apart from the CH and ON subpopulations described above, the CN of the full AN-CN-IC-A1 model also contained 1000 primary-like (PL) neurons. PL neurons make up approximately 47% of the ventral CN in the brain [1], suggesting that they might play a significant role in auditory processing. Although their contribution to the preprocessing of AN discharge patterns is perhaps less clear than that of the CH and ON subpopulations, PL cells were included in the model architecture to investigate their effect on auditory class learning. PL cells essentially transcribe AN firing in the brain [1] and were, therefore, modelled using strong 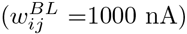 one-to-one AN→PL afferent connections and no inhibition 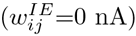 within the PL area. The discharge properties of the model PL neurons were found to correspond closely to those reported experimentally (Fig. 4, left column).

A grid search heuristic was applied to the full AN-CN-IC-A1 model to find the hyper-parameters that produce the best model performance on the two vowel category learning task (see Supplemental Materials for details). Similarly to the reduced AN-A1 model, mutual information was calculated to evaluate the performance of the full AN-CN-IC-A1 model. The following parameter values were found to result in the best model performance: 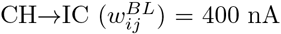, 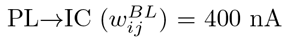 and 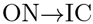 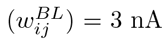 connection magnitudes; the magnitude of the within-IC inhibition 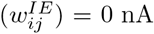; the LTD magnitude of the IC→A1 connections (*α_d_*) = −0.015.

It was found that, unlike the reduced AN-A1 network, a well parameterized full AN-CN-IC-A1 model of the auditory brain was able to solve the two vowel categorization task by developing many A1 neurons with high levels of vowel class identity information approaching the theoretical maximum of 1 bit (Fig. 3, pink). The vowel category information carried in the discharges of the A1 neurons of the full AN-CN-IC-A1 model increased substantially during training (Fig. 3, dotted pink vs continuous pink). Our results, therefore, indicated that the presence of the non-plastic CN micro-architecture converging on the IC indeed helped the plastic A1 learn to produce stimulus class selective responses.

### Generalisation of Learning

We have demonstrated that the trained full AN-CN-IC-A1 model was capable of correctly recognising different exemplars of vowels belonging to either vowel class /i:/ or /a/, despite the high variability even between the input AN spike rasters in response to the different presentations of the same vowel exemplar. It was possible, however, that the model overfit the data and only learnt the particular vowel exemplars presented during training, instead of exploiting the statistical regularities within the stimuli to develop generalised representations of the two vowel classes. To test whether this was the case, we synthesised twelve new exemplars for each of the two vowel classes /i:/ and /a/. The formants of the new vowel stimuli were different to those used in the original stimulus set. Each of the new vowels were presented to the network twenty times. It can be seen in Fig. 3 (green) that many of the A1 cells of the full AN-CN-IC-A1 network trained on the original and tested on the new vowels reached high (up to 0. 92 bits) levels of single cell information about the vowel class identity approaching the theoretical maximum of 1 bit. This suggests that the network indeed learnt general representations of vowel classes /i:/ and /a/, rather than overfitting by learning only the particular vowel exemplars presented during training.

### The Importance of CN and IC Microarchitecture and Connectivity

Having shown that, unlike the reduced AN-A1 model, the full AN-CN-IC-A1 model was capable of learning robust representations of vowel class identities, we investigated next whether the particular microarchitecture and connectivity of the subcortical stages CN and IC was important for the improved AN-CN-IC-A1 model performance, and whether the addition of the two subcortical stages was helpful due to their AN jitter removal properties as hypothesised.

An additional simulation was run to confirm that the particular microarchitecture of the CN and its subsequent convergence on the IC, rather than the pure addition of extra processing layers, improved the performance of the four stage AN-CN-IC-A1 model compared to the two stage AN-A1 model on the vowel class identity learning task. To this accord, a simple four-stage fully connected model lacking the detailed CN and IC microstructure and connectivity (see Fig. 1C, and Materials and Methods for details) was constructed and evaluated using the original two vowel class learning paradigm. Fig. 3 (teal) demonstrates that this simple four-stage network achieved very little information about the identity of the vowel stimuli (no more than 0.28 bits). This suggests that the pure addition of extra processing stages within a spiking neural network model does not help with auditory category learning. Instead, the pre-processing within the particular microarchitecture of the three subpopulations of the CN followed by their convergence on the IC is necessary for such learning to occur.

### The Importance of CN Subpopulations

In order to verify that each of the three CN subpopulations-CH, ON and PL-was important for enabling the full AN-CN-IC-A1 network to learn robust representations of vowel class identities, the performance of the model was evaluated when each of the CN subpopulations was knocked out one by one. Every time one of the CN subpopulations was eliminated from the model, the network parameters were re-optimised to find the best possible classification performance by the new reduced model architecture. Tbl. 2 demonstrates that the removal of any of the three subpopulations of the CN resulted in significantly reduced performance of the AN-CN-IC-A1 model on the vowel class identity recognition task, thus suggesting the importance of all three CN subpopulations in enabling auditory class learning.

### Quantifying Jitter Removal in CN and IC

So far we have demonstrated that the precise microarchitecture and connectivity of the CN and IC are important for enabling the full AN-CN-IC-A1 model to learn robust representations of vowel class identities. Here we test whether the subcortical stages CN and IC indeed remove AN jitter as originally hypothesised. To confirm this, we compared the firing pattern similarity scores between the AN and IC (see Materials and Methods for details). The scores varied between 0 and 1, with high scores indicating high levels of similarity between the corresponding spike rasters and, consequently, low levels of jitter. The high Same Exemplar and Different Exemplars IC scores in Tbl. 1 suggest that the IC firing rasters in response to the different presentations of the same vowel exemplar, or in response to the different exemplars of the same vowel category are highly similar and hence are mostly jitter free. This is in contrast to the corresponding AN scores, which are all significantly lower due to the space and time jitter.

**Table 2.** Maximum single cell information within the output A1 stage of the best performing re-optimised full AN-CN-IC-A1 model when different CN subpopulations of neurons (chopper, onset or primary-like) were selectively knocked out. Y and N indicate that the relevant subpopulation is either present or absent, respectively. The theoretical maximum for the single cell information measure for two auditory classes is 1 bit. The maximum information is only achieved when all three subpopulations are present.

| Chopper | Onset | Primary-Like | A1 Information (bits) |
| --- | --- | --- | --- |
| Y | Y | Y | 1 |
| Y | N | Y | 0.93 |
| Y | Y | N | 0.89 |
| Y | N | N | 0.81 |
| N | Y | Y | 0.36 |
| N | N | Y | 0.18 |
| N | Y | N | 0 |

## Discussion

This work has demonstrated that spike-time jitter inherent to the auditory nerve (AN) firing may prevent auditory category learning in the plastic cortical areas of the auditory brain as evidenced by the poor performance of the reduced AN-A1 or the simple four-stage spiking neural network models of the auditory brain on a controlled and very simple vowel categorisation task. While past research suggested that input spike jitter can be reduced by the intrinsic properties of spiking neural networks [23] and STDP learning [24], such jitter reduction works on the scale of a few milliseconds, rather than tens of milliseconds characteristic of the AN jitter. A jitter removal preprocessing stage is, therefore, important in order to enable the plastic auditory cortex to learn auditory categories. Here we have shown that the ventral cochlear nucleus (CN) followed by the inferior colliculus (IC) are able to do just that. In particular, we have demonstrated that chopper (CH) and onset (ON) subpopulations of the CN and their subsequent convergence on the IC have the right connectivity and response properties to remove space and time jitter in the AN input respectively.

Our simulation results also demonstrated the importance of primary-like (PL) neurons in the CN for enabling the auditory cortex to learn auditory categories. The PL subpopulation simply transcribes AN firing and, therefore, is unlikely to play a role in AN jitter removal. We are, therefore, still unsure what its role in auditory category learning might be. It is possible that PL input is necessary to simply introduce a base level of activation within the IC. Our simulations, nevertheless, have demonstrated that the removal of any of the three subpopulations of the CN (CH, ON or PL) resulted in a significant drop in maximum single cell information within the A1 stage of the trained AN-CN-IC-A1 model (Tbl. 2). This suggests that the AN-CN-IC-A1 model has the minimal sufficient architecture for learning auditory categories.

In this paper we hypothesised that the full AN-CN-IC-A1 model would utilise the Continuous Transformation (CT) learning mechanism to develop stimulus class selective response properties in the A1. For CT learning to be able to drive the development of output neurons that respond selectively to particular vowel classes, the spike rasters in the preceding neuronal stage in response to the different presentations of the same exemplar of a vowel or of different exemplars belonging to the same vowel class must be similar to each other. The presence of spike jitter at any stage of processing will destroy these similarity relations needed for CT learning to operate. The firing pattern similarity scores shown in Tbl. 1 demonstrated that the spike raster similarity/dissimilarity relations required for CT learning to operate were restored in the IC compared to the AN of the full AN-CN-IC-A1 model through the de-jittering preprocessing within the CN and IC. This, in turn, enabled the plastic A1 of the full AN-CN-IC-A1 model to learn vowel categorisation through the CT learning mechanism. The structurally identical A1 layer of the reduced AN-A1 or the simple four-stage models failed to learn from the unprocessed input AN firing patterns due to the space and time jitter breaking the stable AN firing patterns that are necessary for CT learning by STDP to operate.

We hypothesised that space jitter in the AN was removed by CH neurons in the CN, because anatomical studies suggested that CH neurons had the appropriate connectivity from the AN for the task. Similar connectivity, however, is shared by the primary-like with notch (PLn) subpopulation of the CN, suggesting that they may also take part in AN space jitter removal. Neurophysiological evidence, however, suggests that the two cell types have different intrinsic properties [1,25], and the response properties of the CH stage of the AN-CN-IC-A1 model optimised for space jitter removal were found to be more similar to those of the real CH rather than PLn cells (i.e. they do not phase-lock to the stimulus). This suggests that CH cells are more likely to be important for auditory category learning in the brain than PLn neurons.

The simplicity of the synthesised vowel stimuli and the small number of exemplars in each stimulus class are not representative of the rich auditory world that the brain is exposed to during its lifetime. The model, therefore, needs to be tested on higher numbers of stimuli, as well as on more complex and more realistic stimuli, such as naturally spoken whole words, in future simulation work. The two vowel classification problem, nevertheless, was suitable for the purposes of demonstrating the necessity of subcortical pre-processing in the CN and IC for preparing the jittered AN input for auditory category learning in the cortex. The appropriateness of the task is demonstrated by the inability of the reduced AN-A1 and the simple four-stage models of the auditory brain to solve it.

We took inspiration from the known neurophysiology of the auditory brain in order to construct the spiking neural network models described in this paper. As with any model, however, a number of simplifying assumptions had to be made with regards to certain aspects that we believed were not crucial for testing our hypothesis. These simplifications included the lack of superior olivary complex or thalamus in our full AN-CN-IC-A1 model, the nature of implementation of within-layer inhibition in both the AN-A1 and AN-CN-IC-A1 models, and lack of top-down or recurrent connectivity in either model. While we believe that all of these aspects do affect the learning of auditory object categories to some extent, we also believe that their role is not crucial for the task. Therefore, we leave the investigation of these effects for future work.

The full AN-CN-IC-A1 model described in this paper possesses a unique combination of components necessary to simulate the emergent neurodynamics of auditory categorization learning in the brain, such as biologically accurate spiking dynamics of individual neurons, STDP learning, neurophysiologically guided architecture and exposure to realistic speech input. Due to its biological plausibility, the model can be used to make neurophysiologically testable predictions, and thus lead to further insights into the nature of the neural processing of auditory stimuli. For example, one of the proposed future neurophysiological studies would compare the levels of jitter in the real AN and IC in response to the same auditory stimuli, with the expectation being that the level of jitter will be significantly reduced in the IC.

## Materials and Methods

### Stimuli

A stimulus set consisting of twelve exemplars of each of two vowels, /i:/ and /a/, was generated using the Klatt synthesiser [17]. Each 100 ms long sound was created by sampling each of the three vowel formants from a uniform 200 Hz distribution centered around the corresponding formant frequency as reported by [13] for male speakers. The variability in formant frequencies among the twelve stimulus exemplars was consistent with the range of variation present in natural human speech as demonstrated in Fig. 2. Furthermore, informal tests showed that greater variation in vowel formant frequencies resulted in vowel exemplars that sounded perceptually different to /i:/ or /a/. A fundamental frequency (*F*_0_) of 100 Hz was used for all stimuli.

The vowel stimuli belonging to the two classes, /i:/ and /a/, were presented in an interleaved fashion and separated by 100 ms of silence. The silence encouraged the models to learn separate representations of each individual vowel class and to avoid learning any transitions between vowel classes. We used 200 training and twenty testing epochs, whereby each epoch consisted of the first exemplar of vowel /i:/ followed by the first exemplar of vowel /a/, followed by the second exemplar of vowel /i:/ and so on up to the last twelfth exemplar of vowel /a/. Twenty (rather than one) test epochs were used because, due to the stochasticity of AN responses, input AN spike patterns in response to repeated presentations of the same sound were not identical. Informal tests demonstrated that on average the order in which the vowel exemplars were presented did not make a qualitative difference to the performance of the trained models. It did, however, introduce higher trial to trial variability. Hence, we fixed the presentation schedule for the simulations described in this paper for a more fair model comparison.

### Continuous Transformation Learning

The CT learning mechanism was originally developed to account for geometric transform invariance learning in a rate-coded neural network model of visual object recognition in the ventral visual stream [7], but has recently been shown to also work in a spiking neural network model of the ventral visual stream [8]. A more detailed description of CT learning for vision can be found in [26].

In vision, simple changes in the geometry of a scene, such as a shift in location or rotation, can generate a multitude of visual stimuli which are all different views, or “transforms”, of the same object. CT learning was at its origin an attempt to understand how the brain can form representations of visual objects which are not confused by such transformations, i.e. they are “transform invariant”. At first glance it may seem that there is no obvious analogue of such “transformations” in the auditory world. For many classes of natural auditory stimuli, however, their location in “frequency space” depends on the physical characteristics of the sound source. For example, the changes in physical dimensions of the resonators of the vocal tract would create “transformations” of vocalisation sounds. Such changes would happen due to variations in the placement of the tongue or the jaw when the same or different speakers produce the same speech sound. Thus, many natural auditory objects are prone to shifts in frequency space that are not too unlike the shifts in retinotopic space observed when visual objects undergo geometric transformations. We, therefore, propose that CT learning may play a crucial role in auditory category learning.

The original CT learning mechanism relies on the presence of a significant overlap between input representations of temporally static stimulus transforms; in other words, neural representations of “snapshots” of the same object taken from somewhat different points of view often exhibit areas of high correlation which can be discovered and exploited by an associative learning mechanism [7,8]. Unlike snapshots of visual objects, auditory stimuli have an essential temporal structure. In order for CT learning to associate similar temporal presynaptic patterns of firing onto the same output neuron by STDP, it is important that the volley of spikes from the presynaptic neurons arrive at the postsynaptic neuron almost simultaneously [8]. If this is not the case, connections corresponding to the presynaptic spikes that arrive after the postsynaptic neuron fires will get weakened due to the nature of STDP, whereby there is strengthening of connections through long term potentiation (LTP) if the presynaptic spike arrives before the postsynaptic spike and weakening of connections through long term depression (LTD) otherwise, thus preventing effective CT learning of the input patterns.

In order to allow CT learning to work for the temporal auditory stimuli, therefore, a distribution of heterogeneous axonal conduction delays needs to be added to the plastic afferent connections. These axonal delays would transform temporal input sequences into patterns of spikes arriving simultaneously at individual postsynaptic cells. The patterns of coincident spikes received by each postsynaptic cell would depend on the cell’s transformation matrix of axonal delays. If an appropriate delay transformation matrix is applied to the input spike pattern, a subset of postsynaptic neurons will receive synchronized spikes from the subset of input neurons encoding similar exemplars of a particular stimulus class, such as a vowel, thus enabling CT learning. Neurophysiological data collected from different species suggests that cortical axonal connections, including those within the auditory brain, may have conduction delays associated with them of the order of milliseconds to tens of milliseconds [27,28].

It is, therefore, suggested that the CT mechanism can enable a spiking neural network to learn class identities of temporal auditory stimuli if over the whole space of different stimulus exemplars belonging to one class, stimuli that are similar to each other physically also evoke similar spatio-temporal firing patterns (i.e. have “sufficient overlap”). Spatial and temporal jitter, for example in the input auditory nerve (AN), add noise to the spatio-temporal firing patterns, and therefore make responses to similar stimuli more dissimilar, hence preventing effective CT learning without additional preprocessing to reduce such jitter.

### Information Analysis

One common way to quantify learning success is to estimate the mutual information between stimulus category and neural response *I*(*S*; *R*). It is calculated as 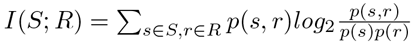, where *S* is the set of all stimuli and *R* is the set of all possible responses, *p*(*S*, *R*) is the joint probability distribution of stimuli and responses, and *p*(*s*) = ∑*_r∈R_p*(*s*,*r*) and *p*(*r*) = ∑_*s∈S*_*p*(*s*,*r*) are the marginal distributions [29]. The upper limit of *I*(*S*; *R*) is given as 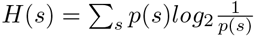, which, given that we had two equiprobable stimulus classes, here equals 1 bit.

Stimulus-response confusion matrices were constructed using a simple binary encoding scheme [12], and used to calculate I(S; R). Binary encoding implies that a cell could either be “on” (if it fired at least once during stimulus presentation), or “off” (if it never fired during stimulus presentation).

We used observed frequencies as estimators for underlying probabilities *p*(*s*), *p*(*r*) and *p*(*s*, *r*), which introduced a positive bias 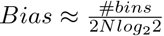, where *#bins* is the number of potentially non-zero cells in the joint probability table, and *N* is the number of recording trials [29]. Given the large value of *N* = 960 in our tests of model performance, the bias was negligible (*Bias* = 0.004 bits) and was ignored.

### Quantifying Spike Raster Similarity

As mentioned above, we hypothesise that a spiking neural network can learn auditory categories through the CT learning mechanism. CT learning relies on a high degree of similarity/overlap between spike rasters in response to different exemplars of one particular stimulus class, such as /i:/ or /a/. Here we describe three indices which quantify the degree of similarity/dissimilarity between spike rasters in response to different presentations of the same exemplar of the same stimulus class (*Same Exemplar Index*), different exemplars of the same stimulus class (*Different Exemplars Index*) or different stimulus classes (*Different Category Index*). Each index varies between 0 and 1, with higher scores indicating a higher degree of similarity between the corresponding firing rasters. Lower scores suggest that the firing rasters being compared are dissimilar due to either the inherent differences between the input stimuli, or due to the presence of spike time jitter that diminishes the otherwise high similarity between the firing rasters being compared.

#### Same Exemplar Index

The *Same Exemplar* (*SE*) index quantifies the degree of similarity between the firing rasters within a particular area (such as AN or IC) in response to different presentations of the same exemplar of a stimulus. Broadly, it calculates the average number of identical spikes across the different presentations of each exemplar *e*_*k*(*s*)_ ∈ {e_1(s)_,…, e_12(s)_} of a stimulus *s ∈* {s_1_, s_2_} in proportion to the total number of stimulus exemplar presentations (*n* ∈ [1, *N*], where *N* = 20 testing epochs). For each presentation of each stimulus we, therefore, constructed a *T×J* matrix 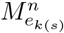 (where *T* = 100 ms is the number of 1 ms time bins spanned by the auditory input, and *J* ∈ {100,1000} neurons is the size of the chosen neural area of the model). Each element *m_tj_* of matrix 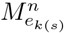 contained the number of spikes produced by the particular neuron *j* ∈ [1, *J*] within the time bin *t* ∈ [1,*T*] in response to the stimulus exemplar *e*_*k*(*s*)_.

If the firing rasters of the chosen area of the model in response to the different presentations *n* ∈ [1, *N*] of the same stimulus exemplar *e*_*k*_(*s*) are similar to each other, then the same slots of the firing pattern matrices 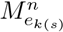 should be non-zero for different *n* ∈ [1, *N*]. Consequently, the following becomes more likely when the proportion of stimulus presentation epochs *n* for which elements of 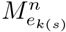 are non-zero across the different presentations of the same stimulus exemplar becomes large: 1) the firing responses within the model area are more likely to be similar; 2) it is likely that less jitter is present in the chosen area of the model; 3) CT learning is more likely to enable postsynaptic cells to learn that the similar, stable, jitter-less responses within the model area belong to the same stimulus class.

We, therefore, computed the matrix 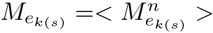, where < > signifies the mean over all the presentation epochs *n* ∈ [1, *N*], and then identified the mean *μ_e_k(s)__* of the hundred largest elements of *M*_*e*_*k*(*s*)__. These were used to compute the final *SE*_*s*_ score for each stimulus *s* ∈ {*s_1_, s_2_*} as *SE*_*s*_ =< *μ*_*e*_*k*(*s*)__ >, where < > signifies the mean over all exemplars *e*_*k*(*s*)_ of stimulus s. A higher *SE_s_* index points to more similarity between the chosen firing rasters in response to the different presentations of the same exemplar of stimulus *s*. Consequently, this also signifies lower levels of jitter present within the layer, since high levels of jitter would disrupt the similarity in firing patterns and result in a lower *SE_s_* index.

#### Different Exemplars Index

The *Different Exemplars* (DE) index quantifies the similarity of the firing rasters within a chosen neural area of the model in response to the different exemplars of the same stimulus class. It is somewhat similar to the *SE_s_* index described above, however, instead of comparing the firing matrices across the different presentations *n* of the same stimulus exemplar *e*_*k*(*s*)_, the firing matrices are compared across the different exemplars *e*_*k*(*s*)_ of each stimulus class *s* ∈ {*s*_1_, *s*_2_}. Consequently, firing raster matrices 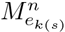 were calculated once again, but this time the average was taken over all the different exemplars *e*_*k*(*s*)_ ∈ {*e*_1(*s*)_, …,*e*_12_(_*s*_)} of stimulus *s* ∈ {*s*_1_, *s*_2_}. That is, we computed 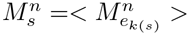, where < > signifies the mean over all the stimulus exemplars. We ^k(s)^ then identified the mean 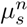 of the hundred largest elements of 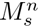 and used them to compute the final *DE_s_* score for each stimulus *s* ∈ {*s*_1_, *s*_2_} as 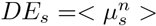, where < > signifies the mean over all *n* ∈ [1, *N*] presentation epochs of each exemplar *e*_*k*_(*s*) of stimulus *s*. A higher *DE_s_* index points to more similarity between the firing rasters within the chosen model neural area in response to the different exemplars *e*_*k*(*s*)_ of stimulus *s*. Consequently, this also signifies lower levels of jitter present within the layer, since high levels of jitter would disrupt the similarity in firing patterns and result in a lower *DE*_s_ index.

#### Different Category Index

The *Different Category* (*DC*) index quantifies the similarity of the different firing rasters within a chosen neural area of the model in response to different stimulus classes. This score is somewhat similar to the *SE*_*s*_ and *DE*_*s*_ scores described above, however, here the rasters are compared across the different stimulus categories *s* ∈ {*s*_1_, *s*_2_}. To this accord, firing raster matrices 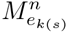 were calculated once again, but this time the average 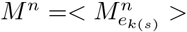 was taken over all the different exemplars *e*_*k*(*s*)_ ∈ {*e*_1_(*s*)__,…, *e*_12(*s*)_} and over all the stimuli *s* ∈ {*s*_1_, *s*_2_}. We then identified the mean *μ^n^* of the hundred largest elements of each matrix *M^n^* and used them to compute the final *DC* score as *DC* =< *μ^n^* >, where < > signifies the mean over all *n* ∈ [1, *N*] presentation epochs. A lower *DC* index points to more differences between the chosen firing rasters in response to the different stimulus categories *s*.

### Spiking Neural Network Models

#### Neuron Model

Apart from the AN, all other cells used in this paper were modelled according to the spiking neuron model by [16]. The model by [16] was chosen because it combines much of the biological realism of the Hodgkin-Huxley model with the computational efficiency of integrate-and-fire neurons. We implemented our models using the Brian simulator with a 0.1 ms simulation time step [30]. The native Brian exponential STDP learning rule with nearest mixed weight update paradigm was used [30]. A range of conduction delays between layers is a key feature of our models. In real brains, these delays might be axonal, dendritic, synaptic or due to indirect connections, but in the model, for simplicity, all delays were implemented as axonal. The [0, 50] ms range was chosen to approximately match the range reported by [31].

**Excitatory Cells:** Neurophysiological evidence suggests that many neurons in the subcortical auditory brain have high spiking thresholds and short temporal integration windows, thus acting more like coincidence detectors than rate integrators [32, 33]. This is similar to the behaviour of Izhikevich’s “Class 1” neurons [16]. All subcortical (CN, IC) excitatory cells were, therefore, implemented as Class 1. To take into account the tendency of neurons in the auditory cortex to show strong adaptation under continuous stimulation [34] Izhikevich’s Spike Frequency Adaptation neurons were chosen to model the excitatory cells in the auditory cortex (A1).

**Inhibitory Cells:** Since inhibitory interneurons are known to be common in most areas of the auditory brain [34, 35] except the AN, each stage of the models apart from the AN contained both excitatory and inhibitory neurons. Inhibitory cells were implemented as Izhikevich’s Phasic Bursting neurons [16]. Sparse connectivity between excitatory to inhibitory cells within a model area was modelled using strong one-to-one connections from each excitatory cell to an inhibitory partner. Each inhibitory cell, in turn, was fully connected to all excitatory cells. Such inhibition implemented dynamic and tightly balanced inhibition as described in [36], which resulted in competition between excitatory neurons, and also provided negative feedback to regulate the total level of firing within an area. Informal tests demonstrated that the exact implementation of within-layer inhibition did not have a significant impact on the results presented in this paper, as long as the implementation still achieved an appropriate level of within-layer competition and activity modulation.

#### Reduced AN-A1 Model Architecture

The reduced AN-A1 spiking neural network model of the auditory brain consisted of two fully connected stages of spiking neurons, the AN (input) and the A1 (output) (Fig. 1B). The AN consisted of 1000 medium spontaneous rate neurons modeled by [11] with CFs between 300-3500 Hz spaced logarithmically, and with a 60 dB threshold. The firing characteristics of the model AN cells were tested and found to replicate reasonably accurately the responses of real AN neurons recorded in neurophysiology studies.

The AN and A1 stages were fully connected using feedforward connections modifiable through spike-time dependent plasticity (STDP) learning. The connections were initialised with a uniform distribution of axonal delays (Δ_*ij*_) between 0 and 50 ms. The randomly chosen axonal delay matrix was fixed for all simulations described in this paper to remove the confounding effect of different delay initialisation values on learning. Informal testing demonstrated that the choice of the axonal delay matrix did not qualitatively affect the simulation results. The initial afferent connection strengths 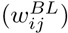 were randomly initialised using values drawn from a uniform distribution. A grid search heuristic was used to find the optimal model hyperparameters (see Tbl. S1 in Supplemental Materials for full model parameters).

#### Full AN-CN-IC-A1 Model Architecture

The full AN-CN-IC-A1 spiking neural network model of the auditory brain consisted of four stages of spiking neurons as shown in Fig. 1A. In contrast to the reduced AN-A1 network, the full AN-CN-IC-A1 model included two intermediate stages between the input AN and output A1 stages to remove time and space jitter present in the AN. These intermediate stages were the CN with CH, ON and PL subpopulations, and the convergent IC stage. The architecture of the three subpopulations of the CN and their corresponding connectivity from the AN is discussed in the Results section. The CN→IC connectivity was the following: CH had Gaussian topological connectivity, whereby each cell in the IC received afferents from a small tonotopic region of the CH subpopulation (*σ* = 2 cells); PL→IC connections were set as one-to-one; ON→IC connections were set up using full connectivity. The AN and A1 stages of the full AN-CN-IC-A1 model were equivalent to those in the AN-A1 model. The IC→A1 connections in the full AN-CN-IC-A1 model were set up equivalently to the AN→A1 connections of the reduced AN-A1 model. Full model parameters can be found in Tbl. S2 (Supplemental Materials).

#### Simple Four-Stage Model Architecture

The simple fully-connected feedforward four-stage model was initialised with randomly distributed synaptic weights 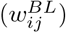 and axonal delays (Δ_*ij*_) between each of the stages, and with STDP learning for the Stage 3→A1 connections (see Fig. 1C). The magnitudes of the feedforward connections 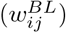 were chosen to ensure that the rate of firing in Stage 3 was similar to that of the equivalent IC stage of the AN-CN-IC-A1 model (approximately 9 Hz). The STDP parameters of the Stage 3→A1 connections were set to the mean of the corresponding optimal values found through the respective parameter searches for models AN-A1 and AN-CN-IC-A1. Full model parameters can be found in Tbl. S3 (Supplemental Materials).

## Acknowledgments

This research was funded by a BBSRC CASE studentship award (No. BB/H016287/1), and by the Oxford Foundation for Theoretical Neuroscience and Artificial Intelligence.

